# Genomic Analysis of Human-infecting *Leptospira borgpetersenii* isolates in Sri Lanka: expanded PF07598 gene family repertoire, less overall genome reduction than bovine isolates

**DOI:** 10.1101/2024.09.17.613401

**Authors:** Indika Senavirathna, Dinesha Jayasundara, Janith Warnasekara, Suneth Agampodi, Ellie J. Putz, Jarlath E. Nally, Darrell O. Bayles, Reetika Chaurasia, Joseph M. Vinetz

## Abstract

*Leptospira borgpetersenii* commonly causes human leptospirosis, including severe disease. The first published analysis of *L. borgpetersenii*, performed on two strains of serovar Hardjo (L550 and JB197), concluded that the *L. borgpetersenii* genome is in the process of genome decay with functional consequences leading to a more obligately host-dependent life cycle. Yet whole genome analysis has only been carried out on few strains of *L. borgpetersenii*, with limited closed genomes and comprehensive analysis. Herein we report the complete, circularized genomes of seven non-Hardjo *Leptospira borgpetersenii* isolates from human leptospirosis patients in Sri Lanka. These isolates (all ST144) were found to be nearly identical by whole genome analysis; serotyping showed they are a novel serovar. We show that the *L. borgpetersenii* isolated from humans in Sri Lanka are less genomically decayed than previously reported isolates: fewer pseudogenes (N=141) and Insertion Sequence (IS) elements (N=46) compared to N=248, N=270, and N=400 pseudogenes, and N=121 and N=116 IS elements in published *L. borgpetersenii* Hardjo genomes (L550, JB197 and TC112). Compared to previously published *L. borgpetersenii* whole genome analyses showing two to three VM proteins in *L. borgpetersenii* isolates from cattle, rats and humans, we found that all of the human *L. borgpetersenii* isolates from Sri Lanka, including previously reported serovar Piyasena, have 4 encoded VM proteins, one ortholog of *L. interrogans* Copenhageni LIC12339 and 3 orthologs of LIC12844. Our findings of fewer pseudogenes, IS elements and expansion of the LIC12844 homologs of the PF07598 family in these human isolates suggests that this newly identified *L. borgpetersenii* serovar from Sri Lanka has unique pathogenicity. Comparative genome analysis and experimental studies of these *L. borgpetersenii* isolates will enable deeper insights into the molecular and cellular mechanisms of leptospirosis pathogenesis.

**Author Summary:** Leptospirosis is an emerging bacterial zoonosis worldwide. *Leptospira borgpetersenii* predominates as the cause of human leptospirosis in some agricultural contexts. We address here the relatively neglected comparative genome analysis of *L. borgpetersenii*. We show here that *L. borgpetersenii* isolated from humans in Sri Lanka have less genome reduction compared to available cattle isolates and have novel virulence characteristics compared to isolates from other animals including cattle and rats.

## Introduction

Leptospirosis, a globally important but neglected bacterial zoonosis [1–6], is caused by gram-negative spirochetes of the genus, *Leptospira*, and is an emerging zoonotic disease worldwide. Leptospirosis is conservatively estimated to affect approximately 1 million people with ∼60,000 deaths per year [2, 4] with estimated Disability Adjusted Life Years (DALYs) annually, which is on par with cholera, typhoid fever and dengue [1, 4, 7–9]. The estimated number of cases of leptospirosis in humans exceeds an average of 500,000 per year, and the case fatality can be as high as 20% [2, 4, 6]. Leptospirosis incidence is strongly predicted to increase over coming years related to climate change [7, 10–14]. Therefore, cases of leptospirosis are likely to become more common as it has already been recognized as a reemerging infectious disease [2]. Identification and characterization of novel *Leptospira* species, which were discovered recently in both pathogen and intermediate lineages [3, 15], are critical for developing novel diagnostic tools for early detection of the disease, for making timely therapeutic decisions [10, 16–19], and to underpin vaccine development [20, 21].

Whole genome sequencing (WGS) has revolutionized in-depth understanding of infection and pathogenesis of leptospirosis at a molecular level [3, 22–24]. Whole genome analysis of new *Leptospira* isolates from different geographic locations has already advance our understanding of the pathogenic mechanisms [25], which may further facilitate the development of better treatment options [3, 20, 26]. The WGS approach has also become a powerful tool for bacterial strain classification and epidemiological typing [5][27, 28]. Leptospiral genome sequences published to date include at least 654 *Leptospira* sequences with most sequences (49%) belonging to *L. interrogans*, followed by *L. borgpetersenii* (7%), *L. santarosai* (6%), and *L. kirschneri* (5%). The size of these genomes varies from 3.9 to 4.6Mb [7]. This list continues to grow [3, 22].

The first whole genome sequence analysis of *L. borgpetersenii* was published by Bulach *et al.* [29]. A recent study published in 2018 reported the genome of *L. borgpetersenii* strain 4E, a highly virulent isolate obtained from *Mus musculus* in southern Brazil [10]. The above-referenced studies identified a total of 3,469 coding DNA sequences (CDSs), 37 transfer-RNAs (tRNAs), 4 ribosomal RNAs (rRNAs), one transfer-messenger RNA (tmRNA) and five riboswitch *loci* in *L. borgpetersenii*. Nevertheless, a fully closed complete genome of *L. borgpetersenii* was reported for the first time based on the genome of laboratory-maintained reference strain, *L. borgpetersenii* serogroup Sejroe serovar Ceylonica strain Piyasena isolated in 1964 (from a male patient in Colombo, Sri Lanka). The complete genome sequences of four recent isolates of *L. borgpetersenii* serovar Hardjo designated strains TC112, TC147, TC129, and TC273 were reported to have 3,345-3,495 coding sequences and 397 to 416 pseudo genes [12]. Recently, the PF07598 gene family that encodes the Virulence Modifying Proteins was reported to encode secreted leptospiral exotoxins that may contribute to the pathogenesis of leptospirosis [25]. While four VM proteins were reported in *L. borgpetersenii* serovar Javanica, in contrast, two Hardjo strains have only three VM proteins [30].

In the present study, we performed whole-genome sequencing, *de novo* assembly, structural, and functional annotation of seven pathogenic *L borgpetersenii* isolates recovered from humans in Sri Lanka, tested the proposed genome reduction hypothesis and compared these isolates with others isolated from different mammalian hosts for genomic content of the PF07598 gene family-encoded VM proteins [29].

## Methods

### *Leptospira* strains and genomic DNA extraction

Isolates for this work were obtained from a large study conducted among febrile patients who were clinically classified as ‘probable’ leptospirosis cases, five from the Teaching Hospital Anuradhapura (FMAS_AP2, FMAS_AP3, FMAS_AP4, FMAS_AP8 and FMAS_AP9), and two from the General Hospital Polonnaruwa (FMAS_PN1, FMAS_ PN4) [8, 16, 31, 32]. Details of patient selection and culture isolation are reported in the original papers [8, 16, 31, 32]. These strains were newly isolated from symptomatic patients and had few passages before genomic DNA extraction for WGS. The organisms were first grown in semisolid EMJH media before being sub-cultured in liquid EMJH medium. Cells were harvested in log phase growth, followed by DNA extraction carried out using the gram-negative bacteria protocol from Qiagen’s DNeasy Blood & Tissue Kit including an RNase clean-up step after proteinase K ^+^ buffer ATL incubation [3]. Extracted DNA was quantified using a Qubit 4 fluorometer (ThermoFisher).

### Sample preparation

Genomic DNA (gDNA) size and integrity was assessed by pulsed field gel electrophoresis (PFGE) method before beginning library preparation. Multiplexed PacBio Single Molecule Real-Time (SMRT) bell libraries were prepared from extracted high quality gDNA using the SMRTbell® Express Template Prep Kit 2.0. To prepare 15-kb libraries, 1µg of genomic DNA was sheared using g-tubes™ from Covaris Woburn, MA, USA and AMPure PB Beads(Pacific Bioscience) were used for the concentration of DNA. The DNA was finally repaired by overnight ligation to the overhanging barcoded 8A adapter (Pacific Bioscience). Blue Pippin ™ size selection (Sage Science, Beverly, Massachusetts, USA) of 4 kb or more was performed according to the manufacturer’s instructions. Conditions for annealing the sequencing primer and binding the polymerase to the purified The SMRTbell ™ template was evaluated using a calculator from RS Remote (Pacific Biosciences).

### Whole-Genome Sequencing and assembly

SMRTbell libraries were generated and sequenced on a PacBio RS II system (Maryland Genomics, Institute for Genome Sciences, University of Maryland School of Medicine). A minimum of 800X read coverage was obtained for all seven isolates. Raw read data were preprocessed using an in-house developed quality control pipeline. Genomes were assembled de novo using Canu 2.1 which were then circularized using Circlator[17] (http://sangerpathogens.github.io/circlator). Two overlapping contigs were recovered in all isolates after completion of the workflow. The annotation was completed in all 7 fully closed genomes using NCBI Prokaryotic Genome Annotation Pipeline with default settings.

### Functional annotation and analysis

Genome-level functional annotation was performed using Prokka v1.13.3 (https://github.com/tseemann/prokka) [33] and the RAST server in our seven closed genomes. CRISPRs and Cas regions were predicted by the CRISPR Cas-finder tool (https://crisprcas.i2bc.paris-saclay.fr/CrisprCasFinder/Index). CRISPRs and Cas regions were extracted from annotated data submitted to the RAST server [34]. The Virulence Factor of Bacterial Pathogen Database (VFDB) was used to predict virulence factors in these *Leptospira* genomes [35]. Mobile elements of the seven isolates were identified by screening using tools at http://www.genomicepidemiology.org/services. BLAST search was performed against the IS finder database for the seven genomes at https://isfinder.biotoul.fr [36]. VM proteins were identified by performing a BLAST search (RAST server) against isolates with known VM proteins.

### *In silico* PubMLST, CG View and Multiple genome alignment

Conventional Multi-locus Sequence Typing (MLST) for the seven isolates against the PubMLST database was performed using seven standardized housekeeping genes https://pubmlst.org/leptospira/ [37]. Fully circularized annotated genomes obtained from the RAST server were uploaded to the CGView server [38], an interactive comparative genomics tool for circular genomes. For identification and alignment of conserved genomic DNA in the presence of rearrangements and horizontal gene transfer, the software package Mauve (https://darlinglab.org/mauve/mauve.html) was used [39, 40]. For multiple alignments, three of our isolates (FMAS_AP8, FMAS_AP9 and FMAS_PN1), strain Piyasena strain JB 197, and L550 were used.

### Methods to identify PF07598 (VM) protein homolog in animal-infecting strains of *L. borgpetersenii*

Several different approaches were used to identify which (or whether) any of the four Sri Lanka isolate VM homologs (orthologs, paralogs) were present in different strains of *L. borgpetersenii* obtained from animals including serovar Hardjo strains HB203, TC112, TC129, TC147, TC273, serovar Ballum strain LR131, and serovar Tarassovi strain MN900 [30]. In the first approach, the Hidden Markov Model (HMM) for the Conserved Protein Domain Family DUF1561 was obtained from NCBI (https://www.ncbi.nlm.nih.gov/). Currently, there is only one Pfam, PF07598, associated with the this Domain of Unknown Function (DUF) (C.f. <u>Pfam: Family:DUF1561 (PF07598)</u> (xfam.org)). Pfam currently uses 16 species and 83 protein sequences to define DUF1561. The putative protein sequences for each genome were obtained from their respective NCBI annotations. The program hmmscan (http://hmmer.org/) was used to search all the annotated proteins against the DUF1561.hmm model. The hmmscan options “-E 0.001 --domE 0.001” were specified for the searches. The hmmscan reported three proteins meeting these criteria in HB203, TC112, TC129, TC147, and TC273 genomes, four proteins meeting these criteria in the LR131 strain, and two proteins meeting these criteria in the MN900 strain. The second and third searching approaches did not use the NCBI protein annotations. This was done to eliminate the possibility that a homolog could have been missed due to an incorrect or missing protein annotation. For the second approach, the liberal method of searching the translations from every ORF over 50 bp in all six reading frames was utilized. These translations were searched against the DUF1516 HMM as described for the NCBI annotations. In the third method, all four of the Sri Lanka protein sequences were compared by tblastn (default parameters) to the nucleotide sequence of the genomes of all six other *L. borgpetersenii strains*. Any hits with a bitscore > 50 was considered putative positive output. This analysis identified exactly three regions in each of the HB203, TC112, TC129, TC147, and TC273 genomes, and four regions in the LR131 strain, and two regions in the MN900 strain. Looking at the annotations associated with those regions (within each strain) revealed that these were the same three annotations found using method one above. Taken together, this leads us to the conclusion that there are only three coding regions that are homologous to the four Sri Lankan proteins in HB203, TC112, TC129, TC147, and TC273 genomes, and four regions in the LR131 strain, and two regions in the MN900 strain.

## Results

The GC content of the isolates were ranged from 39.36%-39.54% (**Table 1)**.

**Table 1.**
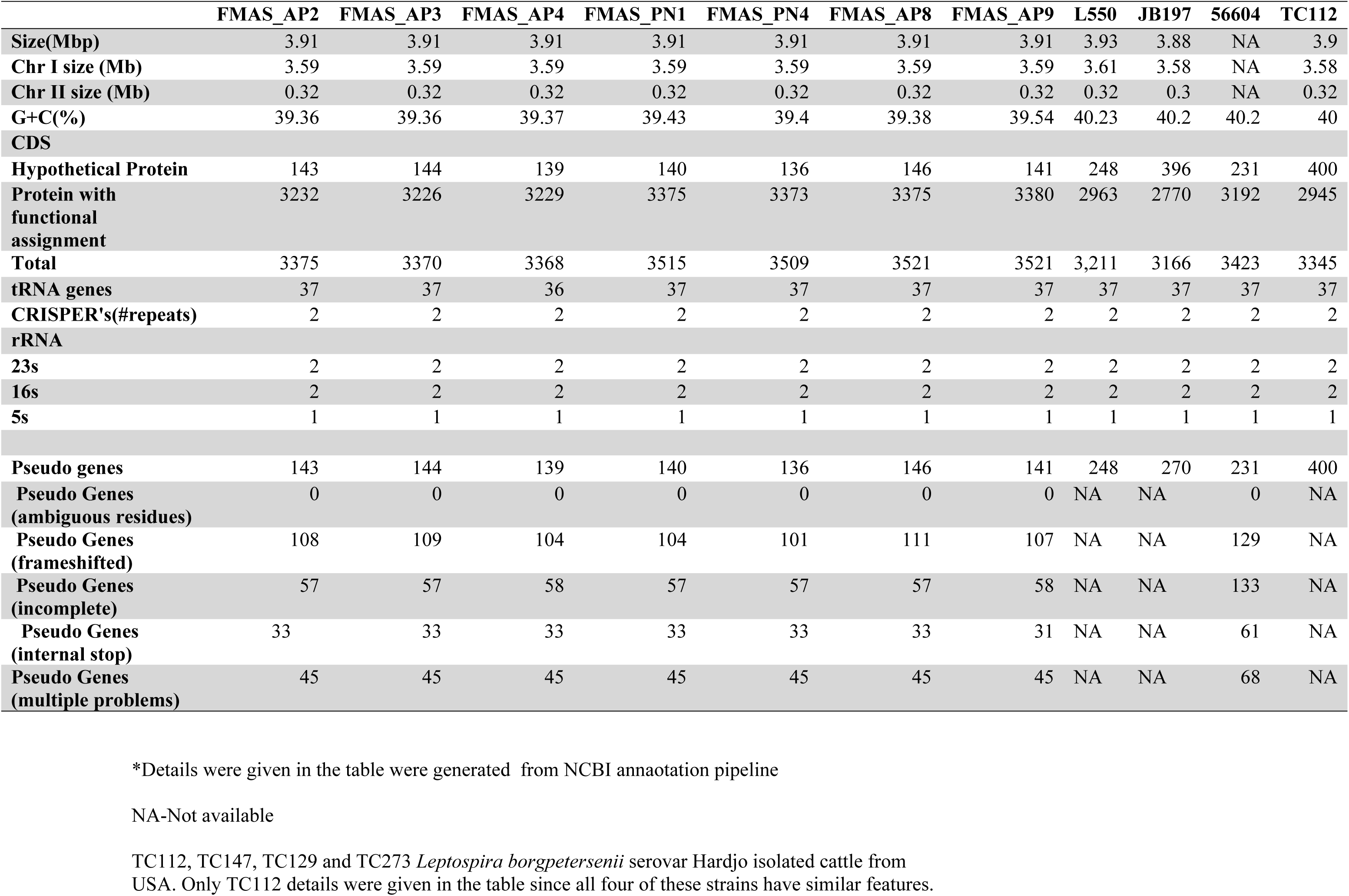
Genome features of Seven *L.borgpetersenii* isolates.

Total coding regions predicted for the isolates ranged from 3368 - 3521. FMAS_AP8 and FMAS_AP9 had same number of coding sequences (CDSs) (3521) while FMAS_AP4 had the lowest number of coding sequences. According to the NCBI annotation, proteins with functional assignment ranged from 3,226 to 3,380 (**Table 1**) while number of hypothetical proteins predicted in the strains had a range of 136-146. FMAS_ PN4 had the lowest number of hypothetical proteins. Two different genomic types were clearly observed based on the coding sequence. FMAS_AP2, FMAS_AP3, and FMAS_AP4 (Group 01) can contain an average of approximately 3,370 protein coding sequences. On the other hand, in FMAS_PN1, FMAS_PN4, FMAS_AP8 and FMAS_AP9 (Group 02) contain about 3,520 protein coding sequences, an increase of about 4.5%. The average protein coding sequences for L550, JB197, 56604, and TC112 are approximately 3,280, representing a 2.7% reduction compared to Group 1 and a 7.3% reduction compared to Group 2. Thirty-seven tRNAs were identified except in FMAS_AP4 in which, only 36 tRNAs were observed. RAST server based subsystem analysis identified 226 in all the strains except in FMAS_AP4 which had only 225. Based on the RAST analysis, CDSs involved in amino acid biosynthesis appeared to be the most abundant subsystem in all strains. The FMAS_AP2 (170) had the highest number of predicted subsystems whereas, strain FMAS_AP4 (168) was predicted to have the least number of subsystems. The subsystem distribution of predicted CDSs in each of the strains is shown in Figure 1.

**Figure 1.**
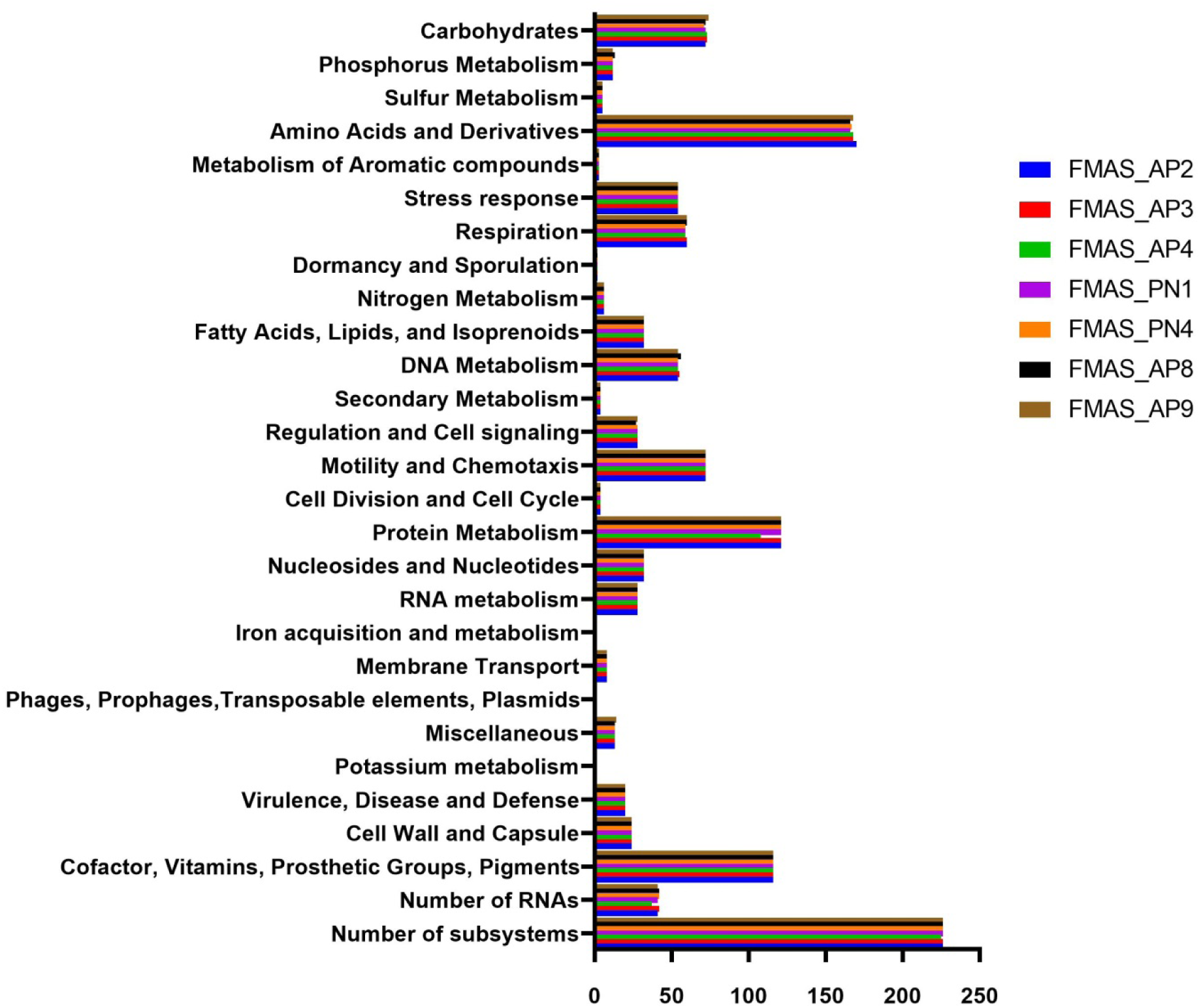
Genomic functional analysis by functional category.

The ST144 MLST profile and CRISPRs and Cas regions predicted by the CRISPR Cas-finder tool and two Crisper-Cas systems were identified in all seven isolates.The circular representation of the seven genomes (CG view) is given in **Figure 1** and the arrangement of the CRISPR system given in **Figure 2**.

**Figure 2.**
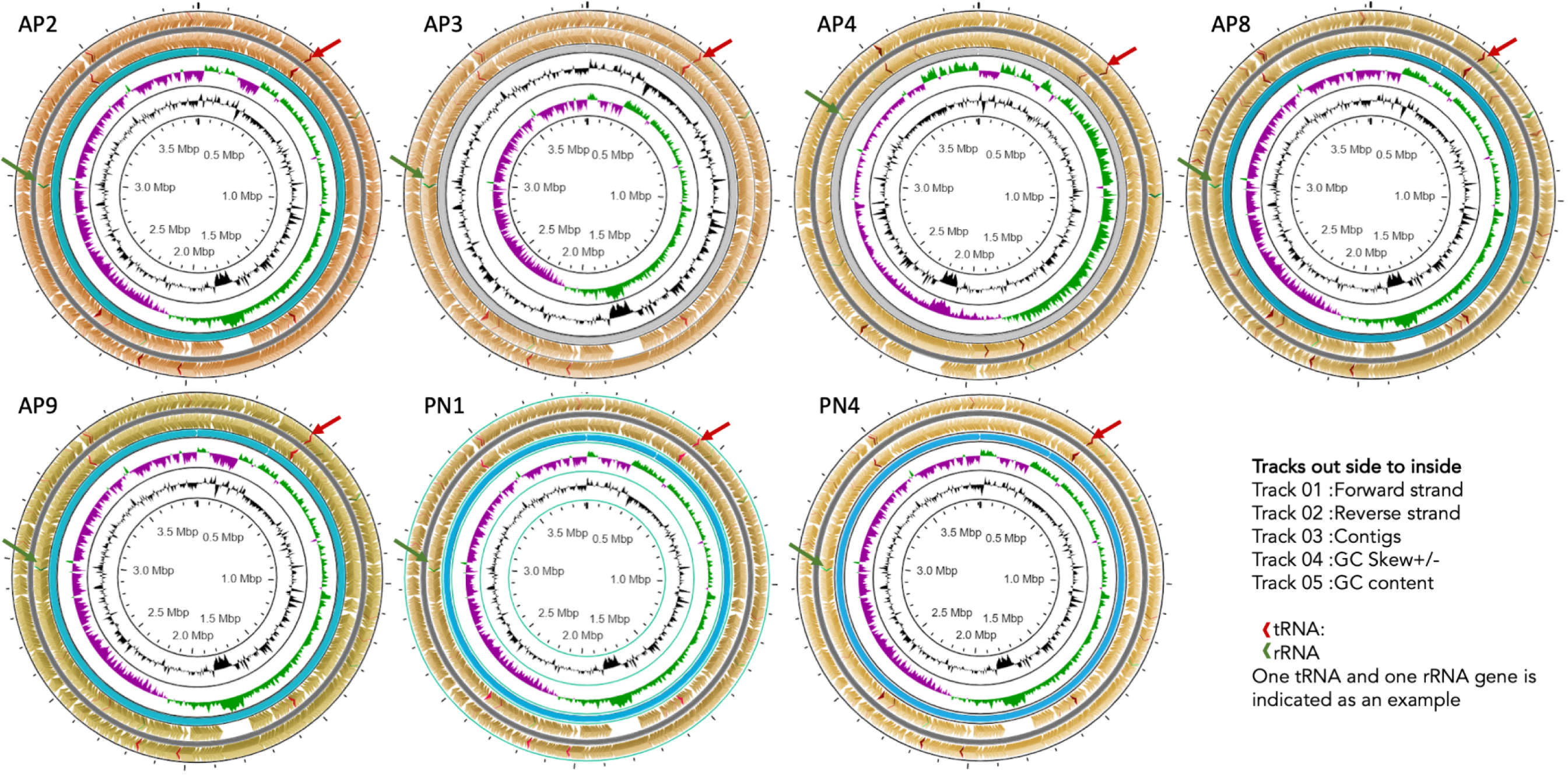
Annotated organization of seven -ex human, Sri Lankan *Leptospira borgpetersenii* genomes using Circular Genome View Plot.

Several putative virulence factors were identified in these *Leptospira* genomes using the VFDB database. Twelve virulence factors were identified in each of the seven isolates (**Table 2).** Five main IS elements were identified in seven isolates such as SLbp8, ISLbp4, ISLbp6, IS1533 and ISLbp5 **(Table 3).**

**Table 2.** Virulence factors identified in Seven Sri Lankan *Leptospira borgpetersenii* Isolates.

| Virulence class | Virulence factor | Gene | Length |
| --- | --- | --- | --- |
| **Adherence** | Mannose-sensitive hemagglutinin (MSHA type IV pilus) | pilB | 1674 |
|  | GroEL(Clostridium) | groEL | 1641 |
| **Antiphagocytosis** | Capsular polysaccharide | Capsule 1 | 1023 |
| **Invasion** | Flagella | flhA | 2118 |
|  |  | cheA | 3195 |
|  |  | cheB | 1047 |
| **Enzyme** | Streptococcal enolase(Streptococcus) | eno | 1299 |
| **Secretion system** | VAS type VI secretion system | clpV | 2232 |
|  |  | clpV | 2544 |
| **Lipid and fatty acid metabolism** | Pantothenate synthesis(Mycobacterium) | panD | 351 |
| **Stress adaptation** | Catalases | katA | 1458 |

\*All seven isolates had similar distribution of virulence factors. VF analyzer:automatic pipeline was used for the systematic analysis and was accessed September 1, 2024.

**Table 3.** Mobile Elements.

| **Main Mobile elements from seven isolates Sequences producing significant alignment** | **IS Family** | **Group** | **Organism** |
| --- | --- | --- | --- |
| **ISLbp8** | ISNCY | ISLbi1 | *Leptospira borgpetersenii* |
| **ISLbp4** | IS4 | IS50 | *Leptospira kirschneri* |
| **ISLbp6** | IS630 |  | *Leptospira borgpetersenii* |
| **IS1533** | IS110 | IS1111 | *Leptospira borgpetersenii* |
| **ISLbp5** | IS5 | IS427 | *Leptospira borgpetersenii* |
| ***L. borgpetersenii* serovar Hardjo, strains L550** |  |  |  |
| **Sequences producing significant alignment** | **IS Family** | **Group** | **Organism** |
| **ISLbp2** | IS256 |  | *Leptospira borgpetersenii* |
| **ISLbp1** | IS4 | IS4 | *Leptospira borgpetersenii* |
| **IS1533** | IS110 | IS1111 | *Leptospira borgpetersenii* |
| **ISLbp4** | IS4 | IS50 | *Leptospira kirschneri* |
| **ISLbp3** | IS982 |  | *Leptospira borgpetersenii* |
| **ISLbp8** | ISNCY | ISLbi1 | *Leptospira borgpetersenii* |
| **ISLbp6** | IS630 |  | *Leptospira borgpetersenii* |
| **ISLbp5** | IS5 | IS427 | *Leptospira borgpetersenii* |
| **IS1501** | IS3 | IS3 | *Leptospira interrogans* |
| ***L. borgpetersenii* serovar Hardjo, strains JB197** |  |  |  |
| **Sequences producing significant alignment** | **IS Family** | **Group** | **Organism** |
| **SLbp2** | IS256 |  | *Leptospira borgpetersenii* |
| **ISLbp1** | IS4 | IS4 | *Leptospira borgpetersenii* |
| **IS1533** | IS110 | IS1111 | *Leptospira borgpetersenii* |
| **ISLbp4** | IS4 | IS50 | *Leptospira kirschneri* |
| **ISLbp3** | IS982 |  | *Leptospira borgpetersenii* |
| **ISLbp8** | ISNCY | ISLbi1 | *Leptospira borgpetersenii* |
| **ISLbp6** | IS630 |  | *Leptospira borgpetersenii* |
| **ISLbp5** | IS5 | IS427 | *Leptospira borgpetersenii* |
| **IS1501** | IS3 | IS3 | *Leptospira interrogans* |

To visualize the general organization of the genome and discover potential genome rearrangements among strains, conserved regions were visualized using a Mauve genome aligner (**Figure 3**).

**Figure 3.**
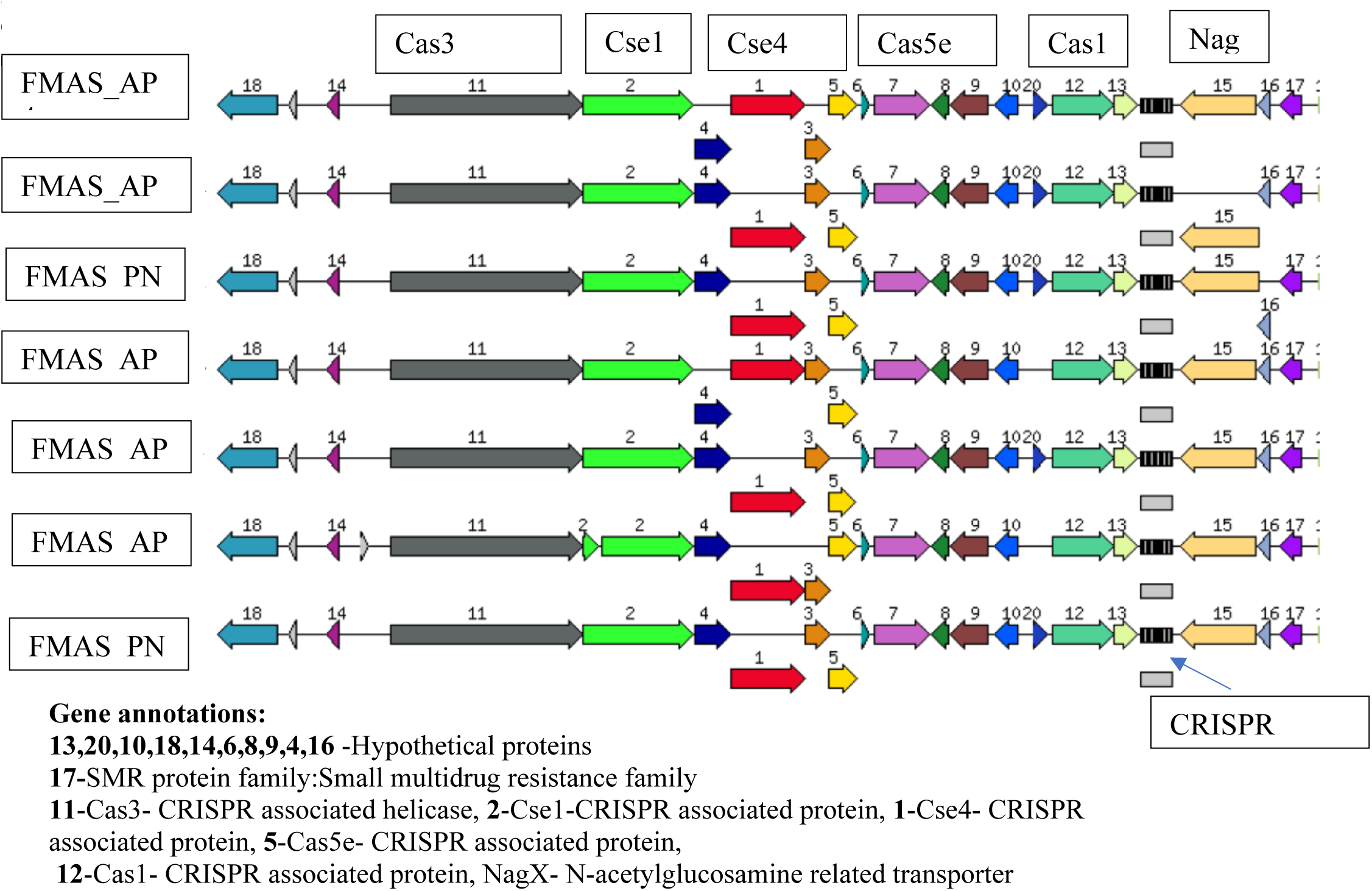
Arrangement of CRISPR/Cas systems in seven -ex human, Sri Lankan *Leptospira borgpetersenii* genomes generated using RAST subsystem analysis.

Large Collinear Blocks (LCBs) were identified. Colored rectangular and variant-specific regions (genomic islands, GI) or white region spaces within or between LCBs were identified in both chromosomes, in all strains. However, chromosome II was highly conserved in all strains (**Fig. 4**). Dimensions and the location of the central LCB on chromosome I was significantly different in our isolates compared to strain piyasena, JB197 and L550. However, FMAS_AP8 and JB 197 had conserved regions throughout the genome. Genomic islands and major genome rearrangements, insertion sequence (IS) elements are often located at the intersection of these rearrangements, which can lead to recombination. The total number of IS elements identified in these strains were 46. The number of pseudogenes identified varied from 136-146. All seven isolates had four VM proteins (**Table 4**).

**Figure 4.**
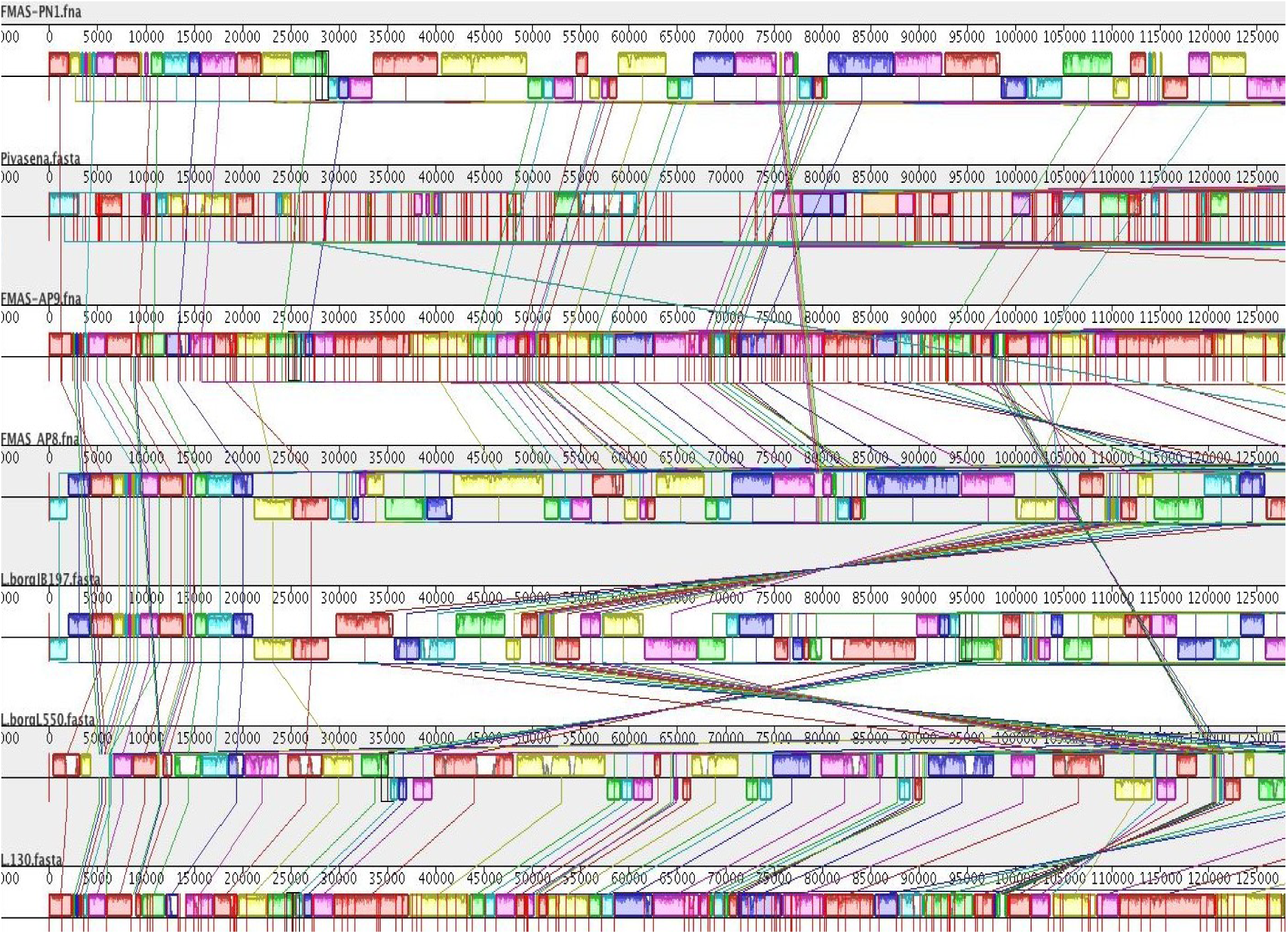
Genomic alignments of seven -ex human, Sri Lankan *Leptospira borgpetersenii* isolates and comparison with reference *Leptospira interrogans* serovar Copenhageni genome using Mauve 2. Snap shot comparing the genomic organization of three Sri Lankan *L. borgpetersenii* isolates with other genomes. Genomes of these strains were aligned and arranged using the Mauve genome aligner in the following order: Top, FMAS_PN1, *L. borgpetersenii* serovar Piyasena, FMAS_AP9, FMAS_AP8, JB 197, L550; at very botton, the reference genome, *L. interrogans* serovar Copenhageni, strain L1-130. Large collinear blocks (LCBs) correspond mainly to conserved syntenic regions, as represented by colored boxes. The lines between the genomes connect the blocks that are conserved between two strains and larger scale rearrangements.

**Table 4.** Comparison of VM Protein Profile Among Seven Ex-human Sri Lankan *Leptospira borgpetersenii* New Isolates and Historical Data.

| Isolate | Species | Serovar | No. VM Proteins | Host |
| --- | --- | --- | --- | --- |
| FMAS_AP2 | *L. borgpetersenii* | No agglutination | 4 | Human |
| FMAS_AP3 | *L. borgpetersenii* | No agglutination | 4 | Human |
| FMAS_AP4 | *L. borgpetersenii* | No agglutination | 4 | Human |
| FMAS_PN1 | *L. borgpetersenii* | No agglutination | 4 | Human |
| FMAS_PN4 | *L. borgpetersenii* | No agglutination | 4 | Human |
| FMAS_AP8 | *L. borgpetersenii* | No agglutination | 4 | Human |
| FMAS_AP9 | *L. borgpetersenii* | No agglutination | 4 | Human |
| Strain 4E | *L. borgpetersenii* | No agglutination | 2 | Rat |
| 55604 | *L. borgpetersenii* | Ballum | 2 | Rat |
| JB197 | *L. borgpetersenii* | Hardjo-Bovis | 3 | Cattle |
| L550 | *L. borgpetersenii* | Hardjo-Bovis | 3 | Human |
| Piyasena | *L. borgpetersenii* | Ceylonica | 4 | Human |
| L49 | *L. borgpetersenii* | Hardjo-Bovis | 3 | Cattle |
| 203 | *L. borgpetersenii* | Hardjo-Bovis | 3 | Cattle |

We found that *L. borgpeterseni* has fewer VM proteins than *L. interrogans*, as exemplified by comparison to the *L. interrogans* serovar Copenhageni str. Fiocruz L1-130 reference genome [41]. The PF07598 gene family encodes a newly identified leptospiral virulence factor family, the Virulence Modifying (VM) proteins. There are four encoded VM proteins, one that is an ortholog of LIC12339 and three that are orthologs of LIC12844. The sequence similarity ranged from 68.34% to 71.24%. The coding region encodes for 638, 632, 629, and 536 amino acids, respectively **(Table 5).**

**Table 5.**
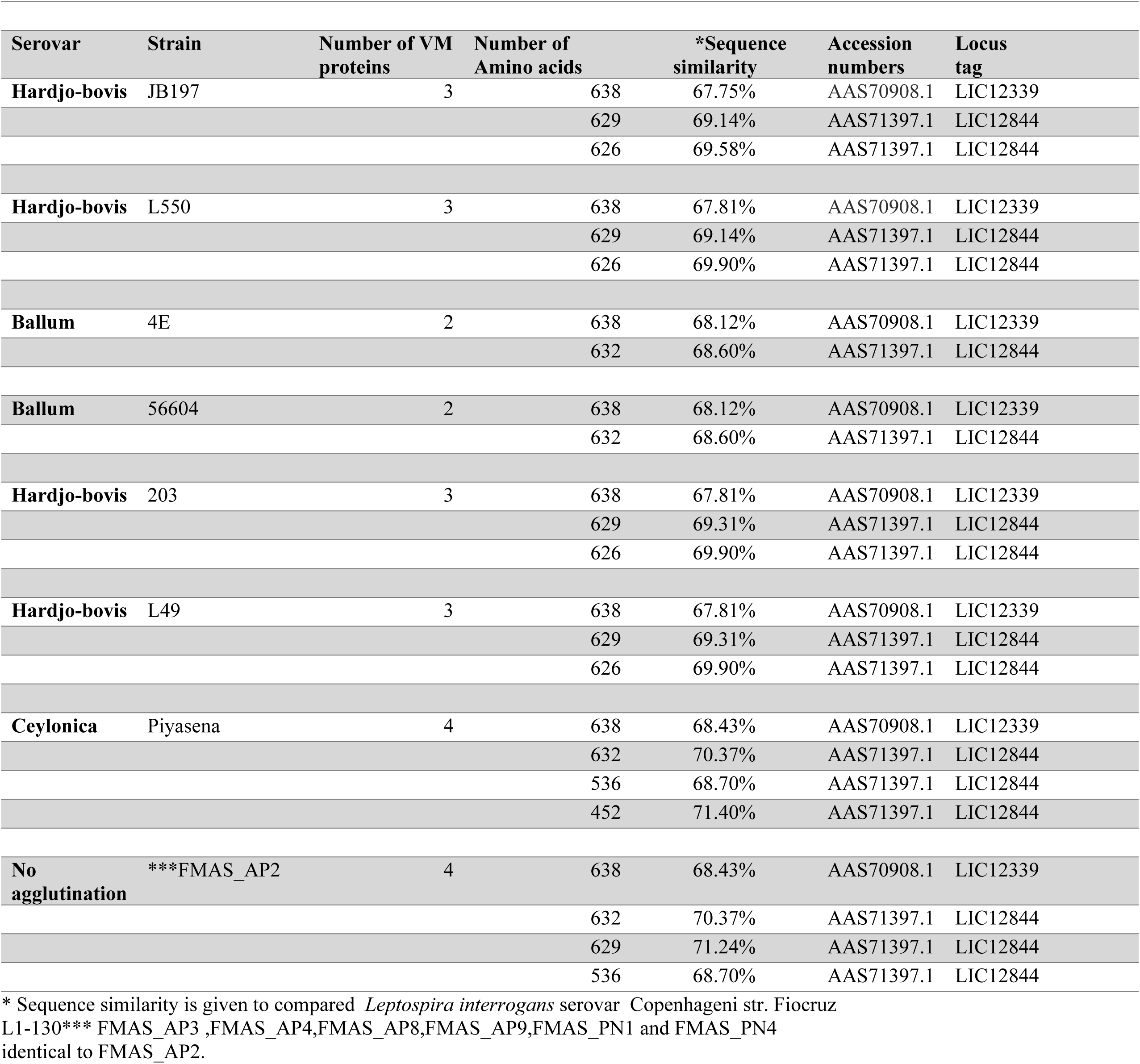
Comparison of the PF07598 (VM Protein) Homologs (Orthologs, Paralogs) in the New Seven Ex-Human *Leptospira borgpetersenii* New Isolates and Historical Sri Lankan Isolates to Ex-Animal Isolates.

We investigated whether the new Sri Lankan isolates shared VM homologs with strains of *L. borgpetersenii* isolated from animal hosts, including bovine isolates of serovar Hardjo strains HB203, TC112, TC129, TC147, TC273, a rodent isolate of serogroup Ballum strain LR131, and a bovine isolate of serovar Tarassovi strain MN900. Three different methods were utilized to identify VM homologs as described, including an hmmscan search of annotated protein sequences of the DUF1561 protein family, searching translations from all ORFs over 50 bp against the DUF1561 HMM, and using tblastn to search the genomes’ nucleotide sequences for any high scoring pairs returned from querying with the known VM protein sequences. Compared to the four VM proteins present in the Sri Lankan strains, collectively, all methods describe the presence of three VM homologs in the serovar Hardjo strains (HB203, TC112, TC129, TC147, and TC273), four VM homologs in the rodent serogroup Ballum strain LR131, and only two VM homologs in the serovar Tarassovi strain MN900.

## Discussion

Here we present the whole genome analysis of new isolates of a novel, non-serotypable *Leptospira borgpetersenii* isolated from humans in Sri Lanka. The main findings are that 1) these isolates from humans, are essentially genomically identical; 2) the level of genome reduction appears to be substantially less than originally proposed for reference *L. borgpetersenii* Hardjo-Bovis strains [29] and therefore these new ex-human Sri Lanka isolates are not simply genomically degenerated parasitic bacteria; and 3) the genomic analysis reflects emergence of a predominant leptospiral strain (ST144), with Sri Lankan bovines as the likely source of human infection.

An increased incidence of human leptospirosis due to *L. borgpetersenii* has been reported worldwide. In a study carried out in the Caribbean archipelago of Guadeloupe during an outbreak, the isolates showed the emergence of the Ballum serogroup (*L. borgpetersenii*), serogroup Icterohaemorrhagiae (*L. interrogans*) [8]. Another report from Malaysia also identified *L. borgpetersenii* serovar Bataviae transmitted by two dominant rat species, *Rattus rattus* and *R. norvegicus* [2]. *L. borgpetersenii* has been reported to cause severe human disease [42–44].

Previous studies have reported genome reduction in *L. borgpetersenii* serovar Hardjo strains L550, JB197 and *L. borgpetersenii* serogroup Ballum serovar Ballum strain 56604 [5,9]. These studies drew a general conclusion that the leptospiral species, *L. borgpetersenii*, has undergone IS-mediated genome shrinkage due to inter-host transmission (not requiring environmental mediated transmission). IS elements are also thought to be important features of the *L. borgpetersenii* genome and mechanisms of genomic decay, contributing to multiple chromosomal rearrangements and pseudogene formation. The total number of coding sequences reported in the three strains were serovar Ballum 56604 (N=2618), serovar Hardjo strains L550 (N=2832) and serovar Hardjo strains JB197 (N=2770) [5,9].

All seven isolates reported in this study belongs to MLST sequence type 144. In the PubMLST database, seven isolates recovered both locally and globally have already been listed under this ST. The first one was the *L. borgpetersenii* serovar Ceylonica isolated from a human in 1964 from Sri Lanka. Other local isolates include human samples from Gampaha, Giradurukotte, Bogammana and a rodent isolate from a black rat in Sri Lanka [32]. The other two are global isolates each from Thailand and Laos. Since all seven isolates from the present study isolated from the dry zone belong to the same ST 144, it might have emerged as the predominant sequence type in that particular geographical region. cgMLST of these seven isolates revealed their clonal group as 267. However, cgMLST data for the previous seven isolates aren’t available for more comprehensive analysis. According to the Mauve alignment genome, strain Piyasena (a previous Sri Lankan isolate) is significantly different from our isolates. FMAS_AP8 and JB 197 had the significant number of conserved regions. The pathogenesis of *L. borgpetereseni* strain Hardjo JB197 is an anomaly [45]; as a laboratory isolate obtained from cattle, this strain is fairly unique for causing acute, lethal disease in hamsters; its chromosome is rearranged significicantly compared to other Hardjo-Bovis strains. However, we have not found this rearrangement of Chromosome II in our *L. borgpetereseni* isolates.

Previous studies have reported genome reduction in *L. borgpetersenii* serovar Hardjo strains L550, JB197 and *L. borgpetersenii* serogroup Ballum serovar Ballum strain 56604 [5,9]. These studies drew a general conclusion that the leptospiral species, *L. borgpetersenii*, has undergone IS-mediated genome shrinkage due to inter-host transmission (not requiring environmental mediated transmission). IS elements are also thought to be important features of the *L. borgpetersenii* genome and mechanisms of genomic decay, contributing to multiple chromosomal rearrangements and pseudogene formation. The total number of coding sequences reported in the three strains were serovar Ballum 56604 (N=2,618), serovar Hardjo strains L550 (N=2,832) and serovar Hardjo strains JB197 (N=2,770) [5,9].

Clustered Regularly Interspaced Short Palindromic Repeats (CRISPR)-associated protein systems are found in bacterial genomes, which are important to generate adaptive immunity against invading exogenous genetic elements such as plasmid and phage infection [33,34]. The Cas gene clusters are quite diverse, and they are frequently encoded by a diverse family of proteins with a wide range of functional domains involved in nucleic acid interaction. Two main classes with six types and numerous subtypes were identified in CRISPR Cas systems based on protein families and features of the architecture of cas loci [35]. In pathogenic and intermediate *Leptospira*, three subtypes subtype I-B, subtype I-C and subtype I-E were recognized. CRISPR Cas systems are not present in non-infectious, saprophytic species [35].

CRISPR Cas Finder tool analysis revealed the presence of two CRISPR-Cas systems in all seven isolates of this study. The CRISPR-Cas systems identified in these Sri Lankan isolates closely resembles the sub type 1E with CRISPR array which was previously reported in *Leptospira borgpetersenii* 56604. It was also identified in other group 1 species like *L. alexanderi, L. alstoni* and *L. mayottensis, L. noguchii, L. santarosai, L. weilii* and *L. fainei* [35]. *L. borgpetersenii* serovar Ballum reported to contain three crisper repeats GGTTCAACCCCACGCATGTGGGGAATAGGCT between 2938442–2938534 [34]. In JB197 and L550 these repeats were not detected [34]. In our seven isolates 6-7 repeats were detected. This shows the variability of our strains compared to reported data in global literature. A recent study conducted in Malaysia has shown the presence of 10 to 16 loci with 1 to 13 spacers in the CRISPR arrays in six *L. interrogans* strains [13]. However, the same study suggested further work was needed before making inferences on this observation with relevance to pathogenicity and environmental adaptation of pathogenic *Leptospira*. [12].

The protein secretory systems that export proteins from the cytoplasm in *L. borpetersenii* were found to be Type I and Type II [5]. However, the VF analyzer earch identified the presence of VAS type VI secretion system in all seven of these isolates. The proteins, that were identified as virulent factors were those coding for adherence, anti-phagocytosis, chemotaxis, mortality (invasion), enzyme, lipid and fatty acid metabolism and stress adaptation. These proteins have also been previously reported as virulence factors in other pathogenic *Leptospira* species. However, the number of virulence factors identified in these seven isolates were comparatively low compared to other pathogenic *Leptospira* [7,36].

Mobile elements (IS Elements) insertion can interrupt coding sequences and lead to pseudogene formation in *Leptospira* [4,8]. The number of IS elements varies not just within species but even within serovars. In *L. borgpetersenii,* a total of approximately 54 ISs scattered among chromosomes of strain 56604 have been identified. This includes 31 copies of IS1533, 15 copies of ISLin1, 4 copies of IS1502, 2 copies of IS1500, 1 copy of IS1501 and 1 copy of ISLin2 [9]. Strains L550 and JB197 have been reported to have 121 and 116 IS elements copies, respectively. In contrast, we found a lower number of IS elements in our isolates, N=46, a comparatively low number [37]. In parallel to this observation, a relatively low number of pseudogenes were observed in human-obtained Sri Lankan isolates (136-146) compared to published genomes of cattle-obtained Hardjo-bovis L550, JB197, 56604, and TC112: N=248, 270, 231, and 400 pseudogenes respectively [12,38]. This could be attributed to relatively high number of mobile elements reported in those three strains which may be related to host-pathogen or pathogen-environment interactions. Five types of mobile elements (ISLbp4) belonging to the IS50 family, were identified in the seven Sri Lankan isolates using a web-based mobile element finder. Strains L550 and JB197 were found to have 9 mobile elements types belonging to different IS families. JB197 was isolated from cattle at slaughterhouses in the United States and the L550 strain was isolated from a human with leptospirosis acquired zoonotically from cattle in Australia. The strain 56604 of the serovar Ballum was isolated from a rat in the west region of China. While genome reduction was observed in above strains, which were probably having exclusive host-to-host transmission, our isolates from human cases, among whom the transmission was probably environment-mediated and a lesser degree genome reduction was observed. However this observation needs to be confirmed by further studies targeting animals, humans, and the environment simultaneously [39]. The amplification of VM proteins in all seven isolates isolated from humans compared with other animal-obtained *L. borgpetersenii* isolates which have 2-3 VM proteins may be relevant to mechanisms of human infectivity and pathogenesis. According to the literature, paralogous (PF07598) exists in all group I pathogens and the number ranges from 2 to 12. Some proteins, such as LA1402, LA 0589, have been shown to be upregulated during infection[40]. *L. borgpetersenii* sv. Javanica strain UI09931 was found to have four distinct types of VM orthologs, including LA0591, LA0769, LA0835, and LA1402 [40]. However, only two distinct orthologs LA1402 (N=1) and LA 0589 (N=3) were found in all seven Sri Lankan isolates. The strains JB197, L550, 203, and L49 had identical VM orthologs in a 1:2 ratio to LA1402 and LA 0589. The strain *L. borgpetersenii* sv. Ceylonica strain Piyasena recovered from human subjects in Sri Lanka had similar number of VM proteins as our seven isolates. However, the number of amino acids coded for in one VM protein is relatively low (452) when compared to the seven isolates (536 aa), but genome accuracy still remains to be validated regarding the true VM protein sequences.

While homologs of the four VM proteins found in the Sri Lankan isolates were identified in alternate animal isolates of *L. borgpetersenii*, they were not consistent across serovar with only three ortho/paralogs found in the serovar Hardjo strains (HB203, TC112, TC129, TC147, and TC273), four homologs in the LR131 serovar Ballum strain, and only two in the MN900 serovar Tarassovi strain. Further, while the number of ortho/paralogs may vary between strains, expression patterns of those VM proteins may also vary by strain and between environmental conditions. For instance, recent analysis of the transcriptome of HB203 (causes chronic disease in the hamster model of leptospirosis) and JB197 serovar (causes severe acute) Hardjo strains cultured at 29°C and 37°C, shows that two of the three VM Hardjo homologs were differentially expressed between strains at both 29°C and 37°C; none of the VM proteins were differentially expressed at the transcriptomic level within strains between temperatures under conditions tested, and *ex vivo* analysis or changes in the NaCl concentrartion mimicking *in vivo* conditions may be required to see upregulation [41][26, 46, 47]. It is notable that between strains, VW expression was higher in the severe disease causing JB197 strain compared to the chronic HB203, which broadly suggests VM gene expression may be associated with acute disease presentation in the hamster. In a proteomic data set looking at the highly similar HB203, TC129, and TC273 strains, there is also evidence of strain-to-strain variation for VM proteins[42]. Collectively these results emphasize the need to further characterize expression of these unique proteins and their role in promoting virulence of pathogenic leptospires. These datta provides some indication that there may be less genome reduction and a larger PF07598 gene family repertoire in human-infecting *L. borgpetersenii* strains, independent of serovar.

## Conclusion

We isolated seven essentially identical *L. borgpetersenii* strains from humans with acute febrile over a three year period. We show that a single Sequence Type, ST144, became the dominant strain to cause human infection in the dry zone of Sri Lanka during this period. Genome reduction, described for *L. borgpetersenii* Hardjo strains L550, JB197 and *L. borgpetersenii* serovar Ballum strain 56604, was observed to a lesser degree in these seven Sri Lankan human isolates. Mauve alignment indicates the presence of conserved regions and genome rearrangement within our isolates. VM protein expansion in human-infecting *L. borgpetersenii* in Sri Lanka may contribute to adaptive mechanisms for survival in the environment leading to human infection.

## Author contributions

**Indika Senavirathna:** Conceptualization, Methodology, Software Data curation, Writing-Original draft preparation. **Dinesha Jayasundara**: Data curation, Visualization, Investigation, Writing-Reviewing and Editing. **Janith Warnasekara**: Visualization, Investigation, Editing. **Suneth Agampodi**: Funding acquisition, Supervision, Project Administration, Software, Validation, Writing-Reviewing and Editing. **Ellie J. Putz**: Investigation, Data curation, Methodology, Writing-Reviewing and Editing. **Jarlath E. Nally**: Investigation, Writing-Reviewing and Editing. **Darrell O. Bayles**: Investigation, Data curation, Methodology, Writing-Reviewing and Editing. **Joseph M. Vinetz**: Supervision, Reviewing and Editing, Project administration, funding acquisition.

## Acknowledgments

We would like to thank Dr. Michael Matthias, Mr. S.K. Senevirathna, and Mr. Milinda Perera for technical assistance, and Mr. Shalka Srimantha and Ms. Chamila Kappagoda for culture maintenance and laboratory support.

## Data availability

Annotated assemblies are available in GenBank under accession numbers:

CP072630:CP072631(https://www.ncbi.nlm.nih.gov/nuccore/?term=CP072630:CP072631[accn])

CP072628:CP072629(https://www.ncbi.nlm.nih.gov/nuccore/?term=CP072628:CP072629[accn])

CP072626:CP072627(https://www.ncbi.nlm.nih.gov/nuccore/?term=CP072626:CP072627[accn])

CP072624:CP072625(https://www.ncbi.nlm.nih.gov/nuccore/?term=CP072624:CP072625[accn])

CP072622:CP072623(https://www.ncbi.nlm.nih.gov/nuccore/?term=CP072622:CP072623[accn])

CP072620:CP072621(https://www.ncbi.nlm.nih.gov/nuccore/?term=CP072620:CP072621[accn])

CP072618:CP072619(https://www.ncbi.nlm.nih.gov/nuccore/?term=CP072618:CP072619[accn])

